# Functional deficits of the inhibitory control system in children with ADHD: A task-based fMRI meta-analysis

**DOI:** 10.64898/2026.10.04.756535

**Authors:** Elizabeth Kammer, Shwetha Patil, Helena Huynh, Zhiyao Gao, Weidong Cai

## Abstract

Inhibitory control deficits have long been proposed as a central neurocognitive mechanism underlying childhood Attention Deficit/Hyperactivity Disorder (ADHD). However, neuroimaging findings remain heterogeneous, partly due to methodological variability across developmental samples and experimental paradigms. The present study aimed to characterize the neural architecture of motor response inhibition in typically developing (TD) children and identify reproducible functional abnormalities associated with pediatric ADHD using stringent, developmentally specific inclusion criteria. We conducted two coordinate-based meta-analyses using Activation Likelihood Estimation (ALE). Following a systematic literature search (up to August 2026), we identified 23 task-based fMRI studies reporting inhibitory-control-related activation in TD children (237 foci) and 17 studies comparing children with ADHD and TD peers (114 foci) during the Stop-Signal Task or Go/No-Go Task. \ Statistical significance was determined using cluster-level family-wise error (FWE) correction. In TD children, ALE revealed a robust inhibitory-control network encompassing the bilateral anterior insula (AI), inferior frontal gyrus (IFG), and the midline pre-supplementary motor area (preSMA) and anterior cingulate cortex (ACC). This spatial organization closely resembled the canonical inhibitory-control network consistently observed in adults, suggesting that the core neural architecture supporting motor inhibition is already detectably organized during development. In the between-group analysis, children with ADHD exhibited hypoactivation within the preSMA. Notably, this region demonstrated striking spatial overlap with the preSMA cluster reliably recruited in TD children during motor inhibition. These findings provide convergent support for inhibitory-control models of ADHD and identify altered preSMA recruitment as the key functional abnormalities associated with pediatric ADHD. While the neural infrastructure supporting inhibitory control is broadly established during childhood, children with ADHD exhibit reduced recruitment of a key cortical node within this network during motor inhibition demands.

## Introduction

Childhood Attention-Deficit/Hyperactivity Disorder (ADHD) is a prevalent neurodevelopmental condition affecting 5% to 10% of children globally ^1–3^. Typically diagnosed during middle to late childhood, ADHD is characterized by persistent and developmentally inappropriate behaviors, including motor restlessness, frequent social interruptions, and a consistent failure to follow instructions. These behavioral markers represent the clinical manifestation of the disorder’s core symptomatic domains: inattention and hyperactivity/impulsivity. If left unaddressed, these symptoms frequently lead to a cascade of functional impairments, including academic underachievement, disrupted peer relationships, and heightened familial stress. Elucidating the internal mechanisms that drive these problematic behaviors is essential not only for refining the conceptualization of the disorder but also for the development of targeted, evidence-based intervention strategies.

A prominent theoretical model of ADHD posits that core behavioral symptoms arise from fundamental deficits within the inhibitory control system^4^. Inhibitory control is a critical executive function that regulates an individual’s actions, emotions, and cognitions. Due to its robust psychometric properties and ease of measurement, motor response inhibition has become the most widely investigated component of this system in experimental settings. The Stop-Signal Task (SST) and the Go/No-Go Task (GNGT) are the gold-standard paradigms used to quantify these inhibitory processes. In the SST, the Stop-Signal Reaction Time (SSRT), typically estimated via the Race Model, serves as a latent measure of stopping speed^5, 6^. In the GNGT, the frequency of commission errors serves as the primary indicator of inhibitory failure. According to Barkley’s inhibitory control model, the behavioral dysregulation observed in children with ADHD has been proposed to stem from a primary failure to suppress urges and impulses that are contextually inappropriate or no longer relevant^4^. Meta-analytic evidence consistently demonstrates that children with ADHD exhibit significantly longer SSRTs and higher commission error rates compared to their typically developing (TD) peers ^7^. While these metrics are reliable indicators of deficit, it is worth noting that they generally yield medium effect sizes, suggesting that while inhibitory dysfunction is a hallmark of the disorder, it may be part of a more heterogeneous neuropsychological profile.

The advent of non-invasive neuroimaging techniques has inspired significant interest in the neural architectures supporting cognition and the mechanisms underlying its dysfunction. Decades of neuroimaging research have established that inhibitory control is mediated by a widely distributed network of cortical and subcortical regions. This inhibitory control network primarily includes the anterior insula (AI), inferior frontal gyrus (IFG), middle frontal gyrus (MFG), pre-supplementary motor area (preSMA), supramarginal gyrus (SMG), and the dorsal striatum, with a well-documented functional dominance of the right hemisphere during active response inhibition^8–10^. Experimental paradigms of response inhibition, particularly the SST and GNGT, have been employed to probe neurobiological signatures of ADHD. A vast body of literature reports significant hypoactivation in key regions in inhibitory control, most notably the IFG and preSMA ^11–26^. However, among individuals with ADHD, the evidence base remains strikingly heterogeneous. Several studies report no significant between-group differences, complicating the search for a reliable biomarker ^27–29^.

Previous meta-analyses have attempted to synthesize these findings, generally confirming a consistent under-recruitment of the salience and frontoparietal networks during motor inhibition in ADHD ^30, 31^. However, these analyses often suffer from critical methodological limitations: they frequently pool pediatric and adult cohorts, aggregate across overly broad task categories, or utilize imprecise contrasts of interest derived from paradigms assessing related but distinct cognitive processes, such as interference monitoring or conflict resolution (e.g., incongruent vs. congruent in Stroop or Flanker tasks).

To address these critical gaps, we conducted a rigorous coordinate-based meta-analysis designed to achieve two primary objectives: (1) to delineate the cortical and subcortical architectures consistently engaged during motor response inhibition in pediatric populations, and (2) to characterize the specific functional neurobiological abnormalities present in children with ADHD during such inhibitory demands. Central to our investigation is determining whether the aberrant activation clusters observed in ADHD fall within the normative inhibitory control circuitry defined by TD children. A distinguishing feature of our approach is the implementation of stringent inclusion criteria to ensure the homogeneity and developmental relevance of the synthesized data. Specifically, we required that: (1) group mean age under 18 years old, thereby isolating pediatric-centered mechanisms from adult-oriented data; (2) task-based fMRI investigations utilize either the SST or GNGT, the gold-standard paradigms for motor inhibition; and (3) each study report specific neuroimaging contrasts that probe the inhibitory control process. By applying these rigorous standards, we aim to provide a more precise and reproducible map of the inhibitory control system in childhood ADHD. We hypothesize that children will recruit key nodes in the adult inhibitory control network, including right anterior insula, inferior frontal gyrus and pre-supplementary motor areas, and children with ADHD will exhibit hypoactivation in these key nodes during inhibitory control.

## Method

This meta-analysis was preregistered in PROSPERO (ID: XXXXXXXX) and adhered to the PRISMA reporting guidelines ^32^. A systematic literature search was conducted independently by four researchers (completed on August 31, 2026) using PubMed and Google Scholar, and manually cross-checked for duplication and missing literature.

### Literature Search and Selection Criteria for Meta-analysis of TD children

A systematic literature search was conducted to identify task-based fMRI investigations of inhibitory control in pediatric populations. The search strategy utilized combinations of the following terms: (“children” OR “adolescent”) AND (“response inhibition” OR “inhibitory control”) AND (“fMRI” OR “functional magnetic resonance imaging”). The initial query yielded 4,418 records. Following the removal of duplicate entries, non-peer-reviewed articles, and studies without available English translations, 2,439 unique records were retained for formal screening.

To ensure the homogeneity and developmental relevance of the data, studies were required to meet six strict eligibility criteria. First, the investigation must have utilized task-based fMRI. Second, participants’ mean age must be under 18 years old. Third, the study must have examined TD cohorts or included a healthy TD control group. Fourth, the experimental design was restricted to either the SST or the GNGT to specifically isolate motor response inhibition. Fifth, the analysis must have reported activation contrasts that probed inhibitory processes, such as Stop > Go or No-Go > Go or Stop/NoGo > Rest (or equivalent control conditions). Finally, studies were required to report whole-brain peak coordinates in a standardized stereotactic space, such as the Montreal Neurological Institute (MNI) or Talairach system. For publications meeting all criteria but lacking coordinate data, we contacted the corresponding authors to request the original peak values.

Following full-text review, 23 studies met all inclusion requirements. This cohort consisted of 17 investigations utilizing the GNGT and 6 utilizing the SST.

### Literature Search and Selection Criteria for Meta-analysis of TD vs ADHD children

A systematic literature search was conducted to identify task-based fMRI studies comparing inhibitory control in pediatric ADHD and typically developing (TD) populations. The search strategy utilized the following Boolean string: (“ADHD” OR “Attention-deficit/hyperactivity disorder”) AND (“children” OR “adolescent”) AND (“response inhibition” OR “inhibitory control”) AND (“fMRI” OR “functional magnetic resonance imaging”). The initial query yielded 2,698 records. After removing duplicates, non-peer-reviewed articles, and studies not available in English, 1,592 unique records were retained for formal title and abstract screening.

To ensure methodological consistency and developmental specificity, studies were required to meet seven stringent eligibility criteria for inclusion in the coordinate-based meta-analysis. First, the investigation must have utilized task-based fMRI. Second, participants’ mean age must be under 18 years old. Third, the study design must have included both a diagnosed ADHD group and a TD control group. Fourth, the experimental paradigms were restricted to either the SST or the GNGT, as these are the gold-standard measures of motor response inhibition. Fifth, the analysis must have reported activation contrasts that probed inhibitory processes. Sixth, the study must have provided a direct statistical comparison of activation between the ADHD and TD groups. Finally, investigations were required to report whole-brain peak coordinates in a standardized stereotactic space. For publications meeting all criteria but lacking coordinate data, we attempted to contact the corresponding authors to obtain the relevant peak values for the contrasts of interest.

Following full-text evaluation, 17 studies met all inclusion requirements and were synthesized in the final meta-analysis. This sample comprised 8 investigations utilizing the GNGT and 9 utilizing the SST.

### Meta-analysis

To identify consistent patterns of neurobiological activity during motor response inhibition, we first converted all the coordinates into the same MNI space and then conducted a coordinate-based meta-analysis using the Activation Likelihood Estimation (ALE) algorithm ^33^. The analysis was implemented in Python using NiMARE (v0.11.1) ^34^. The ALE algorithm models the spatial uncertainty of reported foci from each experiment as 3D Gaussian probability distributions. The width of these distributions was determined by the sample size of each study, accounting for the increased spatial variance associated with smaller cohorts ^33^. These probability maps were merged across experiments to create a Modelled Activation map for each study, which were then aggregated to produce a whole-brain ALE map. Statistical significance was determined through a multi-stage thresholding approach designed to optimize anatomical specificity while maintaining rigorous control over false positives. Specifically, the cluster-defining threshold (CDT) approach was implemented ^35^. A standard cluster-forming threshold of p<0.001 was applied to the initial ALE map to identify candidate clusters. To correct for multiple comparisons across the whole brain, we utilized a Monte Carlo permutation procedure with 10,000 iterations and applied a cluster-level Family-Wise Error (FWE) threshold of *p*<0.05 (corresponding to - log_10_(*p*)>1.3).

Because the present meta-analysis included a relatively modest number of studies, we additionally evaluated the robustness and spatial specificity of the findings using a substantially more stringent cluster-defining threshold (CDT p < 0.00001). In meta-analyses with limited experimental samples, broader cluster-forming thresholds may increase susceptibility to diffuse convergence patterns driven by partial spatial overlap across studies. Applying a more conservative CDT reduces the contribution of weakly convergent voxels, limits spatial expansion or merging of adjacent anatomical regions, and preferentially isolates highly reliable and spatially focal convergence signals. This secondary analysis was therefore conducted as a robustness assessment to determine whether the identified activation patterns remained stable under increasingly stringent statistical criteria.

### Assessing study contribution

We utilized the ALEKernel in NiMARE (v0.11.1) to generate Modelled Activation (MA) maps for each individual experiment included in the dataset^34^. The MA maps represent the spatial probability distribution of reported foci for each study, modeled as 3D Gaussian kernels where the width is determined by the experiment’s sample size. The peak MNI coordinates identified in the FWE-corrected hypoactivation map (e.g., preSMA peak at [0, 18, 52]) were transformed into voxel space using the dataset’s affine matrix. We then extracted the MA values for each experiment at these specific voxel indices.

An experiment was defined as “contributing” to the peak if its MA value at the target coordinate was greater than zero. This threshold ensures that the experiment’s reported foci were spatially proximal enough to the peak to be mathematically smoothed into that specific location. By calculating the ratio of contributing studies to the total number of experiments, we quantified the study support (contribution rate) for the primary finding. This procedure allowed for a qualitative assessment of whether the meta-analytic result was driven by a broad consensus of the literature or a smaller subset of high-weight studies.

Because of high spatial convergency between TD children and children with ADHD, we conducted parallel study contribution analyses anchored at the preSMA peaks identified in both the typically developing (TD) activation map and the ADHD hypoactivation map. This comparative approach allowed us to quantify the degree of literature support for the precise anatomical overlap between consistent functional recruitment in TD children and consistent functional failure in the ADHD population.

## Results

### Consistent Neural Substrates of Inhibitory Control in TD Children

We first performed a coordinate-based meta-analysis to identify the neuroanatomical substrates consistently engaged during inhibitory control in TD children. Following a systematic literature search, we identified 23 fMRI studies that reported whole-brain activation coordinates for TD cohorts using standardized inhibitory control paradigms, yielding a total of 237 foci (see **Figure S1** and **Table S1** for study characteristics).

To quantify spatial convergence across studies, we applied Activation Likelihood Estimation (ALE), a voxel-wise meta-analytic technique that detects regions exhibiting greater convergence of reported activation than expected by chance. The ALE analysis revealed a significant prefrontal-cingulate-insular network of activation encompassing the bilateral IFG and AI, midline preSMA and anterior cingulate cortex (ACC), and right MFG (cluster-forming threshold [CDT] p<0.001, cluster-level FWE-corrected p<0.05, **Figure 1a** and **Table 1**).

**Figure 1.**
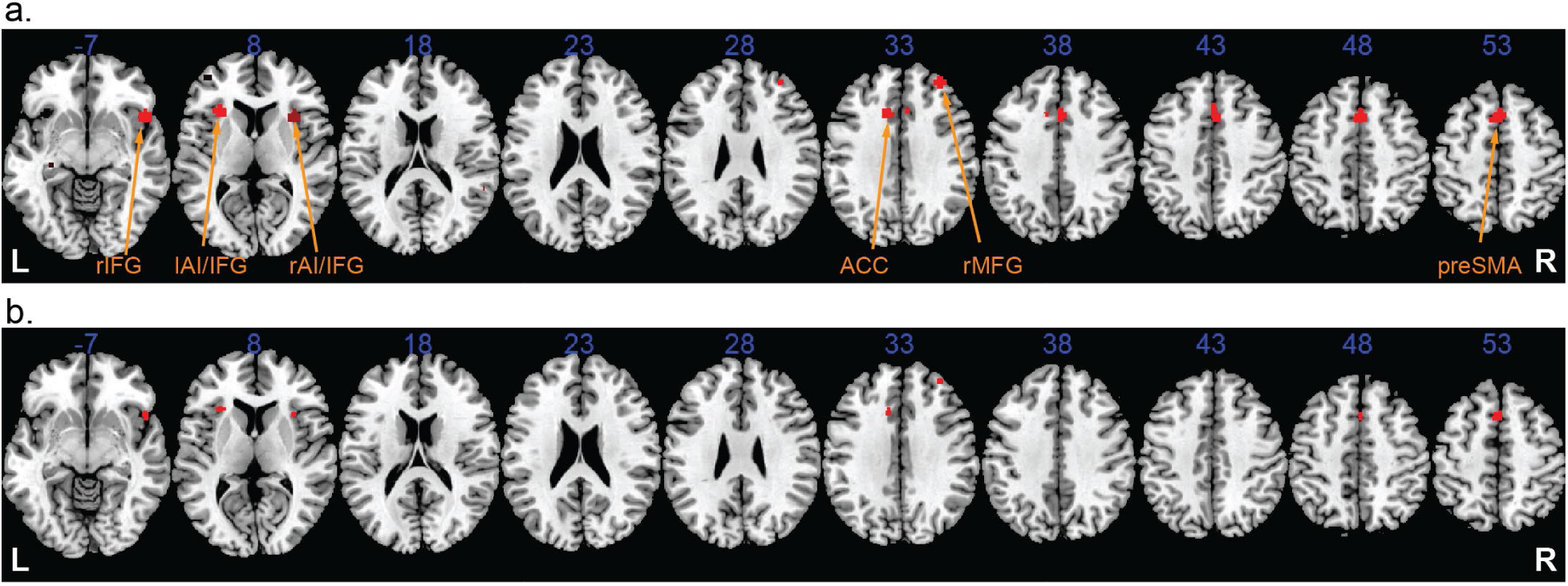
Meta-analysis of neuroimaging studies in TD children. (a) ALE Meta-analysis of 23 inhibitory control neuroimaging studies in children (6 SST, 17 GNGT) revealed consistent activation of the bilateral anterior insula (AI) / inferior frontal gyrus (IFG), anterior cingulate cortex (ACC), pre-supplementary motor area (preSMA), and right middle frontal gyrus (MFG) (CDT p<0.001, cluster-level FWE-corrected p<0.05). (b) A stringent CDT threshold was used (CDT p<0.00001, cluster-level FWE-corrected p<0.05) to further test spatial convergency of significant clusters and demonstrate the robustness of the findings.

**Table 1.** Meta-analytic results in TD children.

| Table 1. Meta-analytic results in TD children |  |  |  |  |  |  |
| --- | --- | --- | --- | --- | --- | --- |
| Threshold: CDT $p < 0.001$, cluster threshold level FWE corrected $p < 0.05$ | | | | | | |
| Cluster ID | Anatomical Label | x | y | z | Stats | Volume (mm3) |
| 1 | preSMA | 2 | 18 | 46 | 4.00 | 2272 |
| 2 | Left AI/IFG | -28 | 22 | 6 | 2.92 | 952 |
| 3 | Right AI/IFG | 50 | 18 | -10 | 2.74 | 1368 |
| 4 | ACC | -10 | 20 | 34 | 1.45 | 576 |
| 5 | Right MFG | 34 | 46 | 32 | 1.38 | 664 |
| Threshold: CDT $p < 0.00001$, cluster threshold level FWE corrected $p < 0.05$ | | | | | | |
| Cluster ID | Anatomical Label | x | y | z | Stats | Volume (mm3) |
| 1 | Left AI/IFG | -30 | 22 | 6 | 3.10 | 296 |
| 2 | preSMA | 0 | 16 | 52 | 2.80 | 272 |
| 3 | Right AI/IFG | 50 | 16 | -8 | 2.42 | 240 |
| 4 | ACC | -10 | 20 | 34 | 1.56 | 128 |
| 5 | Right MFG | 34 | 46 | 32 | 1.56 | 128 |

Stats: $-\log_{10}(p)$ ; AI: anterior insula; IFG: inferior frontal gyrus; preSMA: pre-supplementary motor area; ACC: anterior cingulate cortex; MFG: middle frontal gyrus

Because the present meta-analysis included a relatively limited number of studies (23 experiments), we further evaluated the robustness and spatial specificity of the observed convergence patterns using a substantially more stringent CDT (p < 0.00001). Applying a stricter CDT minimizes the contribution of weakly convergent voxels and preferentially retains regions showing highly consistent and spatially focal activation convergence. This secondary analysis was intended solely as a robustness assessment. Importantly, all principal clusters remained significant following cluster-level FWE correction under this conservative threshold (p < 0.05; **Figure 1b** and **Table 1**), indicating that the identified effects were not dependent on the standard thresholding parameters but instead reflected robust and reliable patterns across studies.

Notably, this spatial distribution closely resembles the cortical inhibitory control system observed in adult neuroimaging studies, which typically centers on the right AI/IFG and preSMA. These findings suggest that the fundamental neural architectures required for inhibitory control are functionally organized in childhood in a manner that resembles mature adult mechanisms.

### Functional Neurobiological Abnormalities in Children with ADHD

Next, we conducted a coordinate-based meta-analysis to identify consistent patterns of functional neurobiological abnormality in children with ADHD relative to their TD peers. Our systematic search identified 17 task-based fMRI studies reporting whole-brain activation differences during inhibitory control tasks, with a total of 114 foci (see **Figure S2** and **Table S2** for study characteristics).

We then employed ALE to determine regions of significant spatial convergence for between-group differences. This initial ALE meta-analysis revealed a trend of diminished activation in children with ADHD within the preSMA relative to TD children (CDT p<0.001, cluster-level FWE-corrected p=0.055, **Figure 2a**).

**Figure 2.**
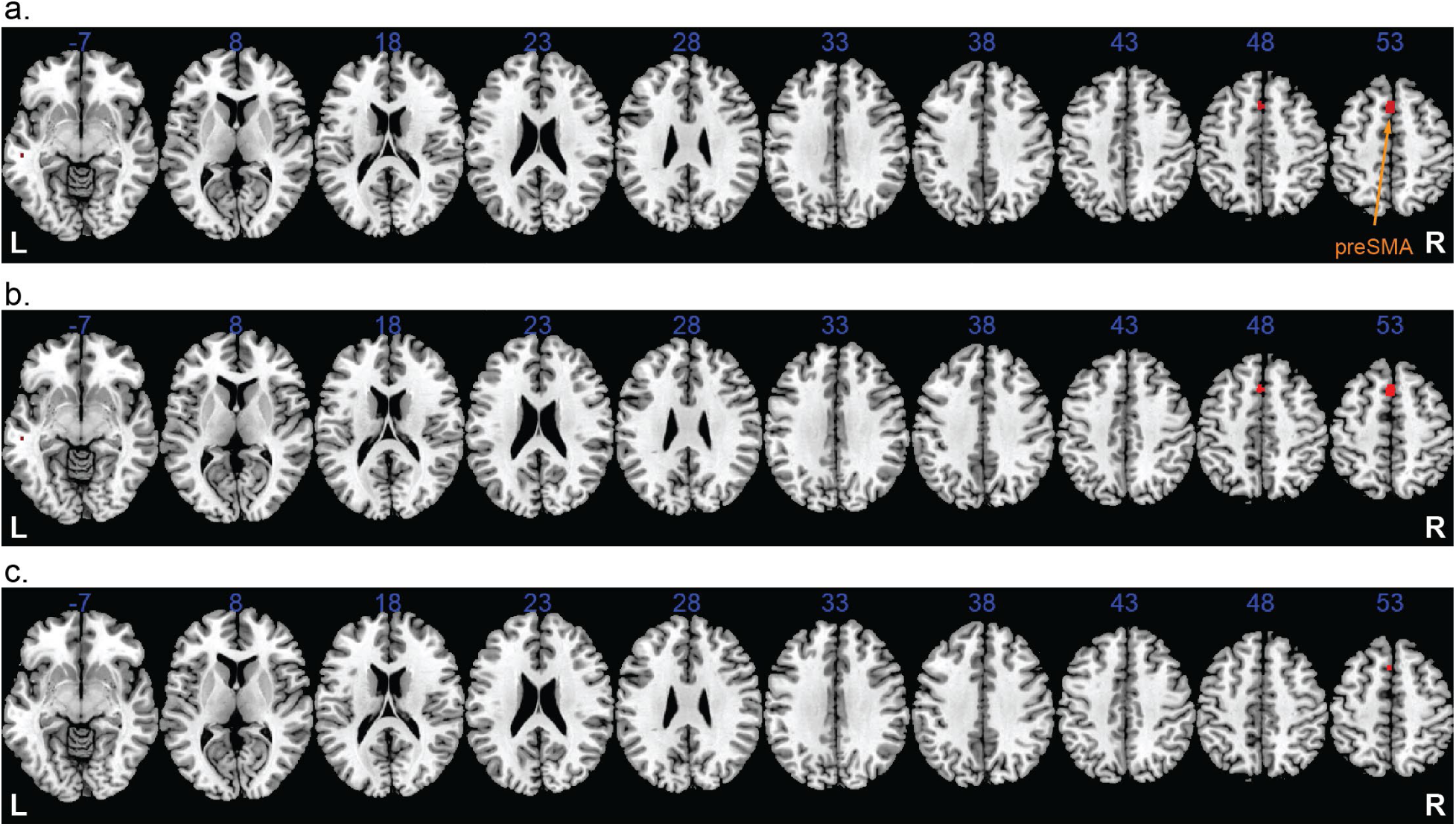
Meta-analysis of neuroimaging studies in TD children versus children with ADHD. (a) ALE Meta-analysis of 17 inhibitory control neuroimaging studies in children (9 SST, 8 GNGT) revealed hypoactivation of the pre-supplementary motor area (preSMA) in children with ADHD, in comparison to TD children (CDT p<0.001, cluster-level FWE-corrected p=0.055). (b) After excluding the study with an exceptionally large cohort ^26^, ALE meta-analysis of 16 studies revealed significantly reduced activation of the preSMA in children with ADHD than TD children (CDT p<0.001, cluster-level FWE-corrected p<0.05). (c) A stringent CDT threshold was used (CDT p<0.00001, cluster-level FWE-corrected p<0.05) to further test spatial convergency of significant clusters and demonstrate the robustness of the findings (all 17 studies included).

To better understand this borderline result, we examined the influence of individual studies on the ALE null distribution. In the ALE framework, spatial uncertainty around each reported focus is modeled as a three-dimensional Gaussian kernel whose width is inversely related to study sample size. Notably, one included study employed an exceptionally large cohort (N = 309) ^26^, substantially exceeding the average sample size of the remaining studies (mean N = 35). Consequently, this study contributed highly focal kernels with unusually narrow spatial uncertainty estimates. Because these high-precision foci showed limited spatial overlap with the broader convergence pattern observed across the remaining experiments, they contributed disproportionately to the maximum ALE values generated during Monte Carlo-based null simulations. This effect inflated the statistical threshold required for cluster-level significance and likely obscured the more consistent convergence signal present across the rest of the literature. To evaluate the robustness of the observed effect, we conducted a sensitivity analysis excluding this disproportionately weighted study. Following its removal, the ALE meta-analysis revealed significant hypoactivation in children with ADHD within the preSMA relative to TD controls (CDT p<0.001, cluster-level FWE-corrected p<0.05, **Figure 2b** and **Table 2**).

**Table 2.** Meta-analytic results in comparing TD and ADHD children (TD > ADHD)

| Table 2. Meta-analytic results in comparing TD and ADHD children (TD > ADHD) |  |  |  |  |  |  |
| --- | --- | --- | --- | --- | --- | --- |
| Threshold: CDT $p < 0.001$, cluster threshold level FWE corrected $p < 0.05$ (16 studies) | | | | | | |
| Cluster ID | Anatomical Label | x | y | z | Stats | Volume (mm<sup>3</sup>) |
| 1 | preSMA | 0 | 18 | 52 | 1.3526 | 616 |
| Threshold: CDT $p < 0.00001$, cluster threshold level FWE corrected $p < 0.05$ (17 studies) | | | | | | |
| Cluster ID | Anatomical Label | x | y | z | Stats | Volume (mm<sup>3</sup>) |
| 1 | preSMA | 0 | 18 | 52 | 1.3915 | 96 |

Stats: $-\log_{10}(p)$ ; preSMA: pre-supplementary motor area

We further assessed the spatial specificity and robustness of this finding using a substantially more stringent CDT (p<0.00001). The preSMA cluster remained significant following cluster-level FWE correction (p<0.05; **Figure 2c** and **Table 2**), even without excluding the study with the exceptionally large cohort. This indicates that the observed hypoactivation reflected a highly consistent and spatially focal convergence pattern rather than a diffuse or threshold-dependent effect.

No significant convergence was observed in subcortical structures (e.g., striatum) or posterior regions at the stringent p<0.05 FWE-corrected level. Furthermore, an analysis of hyperactivation in ADHD yielded no significant clusters, suggesting that functional differences in this population are primarily characterized by a lack of task-relevant engagement rather than compensatory recruitment.

### Spatial Convergence of Normal Activation and Hypo-activation

The preSMA cluster, which exhibited diminished activation in children with ADHD, demonstrated significant spatial overlap with the key region reliably recruited during motor response inhibition in TD children. The Euclidean distance between the peaks of these two clusters was merely 2 mm (**Figure 3**, **Table 1** and **Table 2**). This precise anatomical convergence, achieved despite the stringent p<0.00001 cluster-defining threshold, suggests that altered recruitment is within a central inhibitory control node.

**Figure 3.**
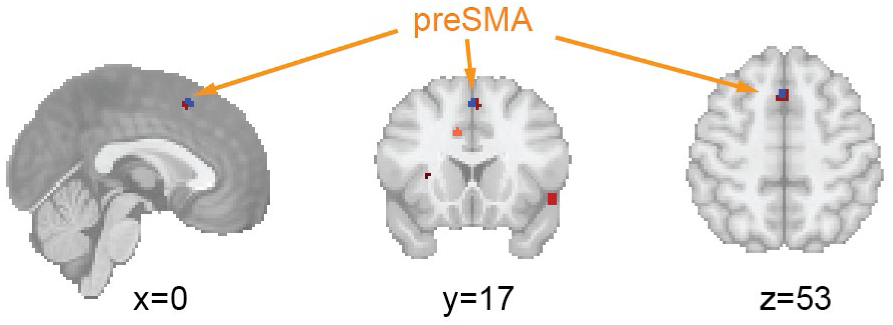
Spatial Convergence of the preSMA activation in TD children (Red) and its hypoactivation in children with ADHD relative to TD children (Blue) (CDT p<0.00001, cluster-level FWE-corrected p<0.05).

### Study Contribution to preSMA Convergence

To evaluate the consistency of the findings across the included literature, we performed a study contribution analysis anchored at the identified preSMA peaks. For studies of TD children, the significant preSMA cluster (Peak MNI: [0, 16, 52]) received direct spatial support from 43.5% of the included experiments (10 out of 23 studies).

In the clinical contrast, the significant cluster of ADHD-related hypoactivation, situated merely 2 mm from the TD peak, was supported by 35.3% of the identified experiments (6 out of 17 studies). Although contribution rates should not be interpreted as direct measures of effect reliability, the comparable degree of study support observed across the TD and ADHD analyses suggests that preSMA hypoactivation during inhibitory control is a recurring, though not universal, finding in the pediatric ADHD literature.

## Discussion

We conducted two rigorous neuroimaging meta-analyses to characterize the neural architecture of motor response inhibition in children and to evaluate the inhibitory control theory of childhood ADHD. Our results demonstrate that children recruit an inhibitory control network that substantially resembles the canonical adult inhibitory control system, specifically involving the AI/IFG and preSMA/ACC, during motor suppression tasks. Notably, compared to TD children, children with ADHD exhibited diminished and inconsistent activation within the preSMA, a critical cortical hub of the inhibitory control system.

### Neural underpinnings of inhibitory control in children

Our coordinate-based meta-analysis provides compelling evidence for a robust and consistent neuroanatomical signature of inhibitory control in TD children. By synthesizing data across 23 fMRI studies using the ALE method, we identified a brain network anchored in the bilateral AI/IFG, and the midline complex of the preSMA and ACC. The emergence of this specific topography is particularly significant as it closely aligns with the core inhibitory infrastructure frequently reported in adults ^8–10^. This spatial correspondence suggests that the fundamental neural architecture required for response inhibition is already detectably organized in childhood^8^. However, inhibitory control function continues to mature through childhood and adolescence^36, 37^. Therefore, our findings imply that the transition from childhood to adulthood in this domain may be characterized more by the refinement and strengthening of existing functional nodes rather than a large-scale reorganization of the inhibitory control system.

### Pediatric neuroimaging findings of inhibitory control deficits in ADHD

Inhibitory control deficits are central to understanding the behavioral problems associated with ADHD ^4, 38, 39^. Over the past two decades, task-based fMRI studies using the SST and GNGT have repeatedly investigated the neural substrates underlying impaired motor inhibition in affected children. Although individual findings have varied considerably across studies, a broad pattern of reduced activation within fronto-striatal and frontoparietal control systems has consistently emerged.

Early investigations primarily implicated right-lateralized prefrontal and cingulate regions during inhibitory control. Initial SST studies reported reduced activation in the right IFG, medial prefrontal cortex, caudate, and ACC in children with ADHD relative to TD controls ^19, 24^. Subsequent event-related and blocked-design paradigms further extended these observations, identifying hypoactivation across the preSMA, MFG, inferior parietal lobule, thalamus, and dorsal striatum during motor inhibition tasks ^12, 13, 17, 22, 40^.

More recent and methodologically rigorous investigations have largely reinforced these earlier findings. Studies employing carefully matched control conditions and larger cohorts have consistently reported diminished recruitment of the IFG/insula, ACC, and preSMA during inhibitory demands in children with ADHD ^14, 15, 26^. Similar patterns have also been observed across medication studies, behavioral intervention paradigms, and task variants probing reactive inhibition and sustained cognitive control ^11, 18, 21, 41, 42^. Collectively, these findings suggest that inhibitory deficits in ADHD are associated with altered engagement of a distributed fronto-cingulo-striatal network responsible for motor suppression and executive control.

Despite this broad convergence, substantial heterogeneity remains across the literature. Several studies have reported weak or absent between-group differences, while the precise anatomical loci of hypoactivation have varied considerably across experiments. This variability likely reflects differences in task design, statistical thresholding, developmental sampling, medication status, and the inclusion of paradigms that recruit partially distinct executive processes beyond motor inhibition itself. These inconsistencies underscore the importance of applying rigorous and highly specific inclusion criteria when attempting to identify reproducible neural abnormalities associated with inhibitory control in pediatric ADHD.

### Meta-analytic research of inhibitory control in ADHD

Several prior meta-analyses have attempted to synthesize the neuroimaging literature on inhibitory control deficits in ADHD, generally supporting the presence of abnormal recruitment within fronto-striatal and salience-related control systems. Early landmark meta-analyses identified reduced activation across the right IFG/AI, Supplementary Motor Area (SMA)/preSMA, ACC, and basal ganglia during inhibitory control tasks in individuals with ADHD ^31, 43^. Similar patterns were later reported in transdiagnostic analyses examining cognitive control across ADHD and Autism, which additionally implicated lateral prefrontal and temporal regions ^44^.

More recent large-scale syntheses have both reinforced and complicated this emerging framework. Zhang et al. ^45^ confirmed consistent hypoactivation within the right IFG/Orbitofrontal Cortex during inhibitory control but also reported regions of increased activation in the SMA, putamen, and temporal cortex, suggesting potential compensatory recruitment mechanisms. In contrast, Samea et al. ^46^ failed to identify significant convergence of inhibitory-control-related abnormalities under stringent cluster-level correction procedures. Together, these discrepancies highlight the substantial methodological heterogeneity characterizing the field.

Several factors likely contribute to these divergent findings. Many previous meta-analyses combined pediatric and adult cohorts, aggregated across broad cognitive-control paradigms, or included contrasts probing related but partially distinct executive processes such as conflict monitoring or interference resolution. Such approaches increase experimental heterogeneity and may obscure the neural architecture specifically associated with motor response inhibition in pediatric ADHD.

The present study was designed to address these limitations by restricting inclusion to pediatric task-based fMRI studies employing the SST or GNGT and contrasts directly isolating inhibitory processes. Under these conditions, our analyses identified preSMA as altered functional recruitment in children with ADHD, suggesting that dysfunction within this central inhibitory-control node represents one of the most stable neurobiological findings across the pediatric literature.

### Hypoactivation of the preSMA during Inhibitory Control in Children with ADHD

Despite the stringent inclusion criteria and the relatively limited sample of eligible studies, our meta-analysis identified the preSMA as a singular, convergent locus of functional deficits in children with ADHD. Specifically, the preSMA was the only region to demonstrate significant hypoactivation during motor response inhibition when compared to TD children.

As a critical cortical hub for action selection and inhibitory control, the preSMA is known to orchestrate the braking signals necessary to halt motor outputs ^47–50^. Neurophysiological and neuroimaging evidence consistently identifies the preSMA as a primary node in the hyperdirect pathway, which allows for the rapid inhibition of the motor system via the subthalamic nucleus ^51–54^. Existing literature has shown that both structural lesions to this area and experimental disturbance of its activity via Transcranial Magnetic Stimulation result in significant impairment of inhibitory performance ^47, 55–57^.

Our findings suggest that preSMA dysfunction may reflect altered engagement of a core inhibitory-control mechanism. Consistent with the meta-analysis finding, a recent study has also found that the preSMA exhibited higher neural instability during inhibitory control in children with ADHD than TD children ^28^. Ultimately, this deficit reflects a compromise in the cortical machinery required for top-down inhibitory control in the ADHD population.

### Consistency of the preSMA deficit

While ALE meta-analyses effectively identify spatial convergence, evaluating the proportion of contributing experiments is essential for understanding literature-wide consensus. The finding that the preSMA node was functionally recruited in 43.5% of the normative TD studies establishes it as a primary locus of developmental inhibitory control. Crucially, the ADHD hypoactivation cluster, situated a mere 2 mm away, was directly supported by 35.3% of the clinical contrasts.

The proportional alignment between normative functional recruitment (43.5%) and clinical hypoactivation (35.3%) is inconsistent with the possibility that the observed convergence was driven solely by a small subset of heavily weighted studies. Instead, it demonstrates that when children with ADHD fail to recruit the necessary cortical engagement for motor suppression, the preSMA is the key locus of that failure.

### Challenges and limitations

As with all coordinate-based meta-analyses, the present findings are constrained by reliance on thresholded peak coordinates reported in published studies. Consequently, the results may be influenced by publication bias, selective reporting practices, and the loss of subthreshold spatial information. A previous meta-analytic attempt failed to identify a significantly consistent neural pattern associated with motor inhibition in children, likely due to the insufficient power of the then-available literature (n=14) ^8^. By expanding to 23 studies, the current analysis successfully delineated a significant inhibitory control network in children that encompasses the cardinal prefrontal, cingulate, and insular regions characteristic of adult inhibitory control topographies. Despite these advancements, the meta-analysis of ADHD-related neural abnormalities remains constrained by a limited sample size. While the identified preSMA cluster is convergent, additional high-powered studies are required to further characterize the full extent of the ADHD-related inhibitory control deficit.

## Conclusion

In summary, our meta-analytic findings demonstrate that while children already exhibit brain activation pattern broadly resembling canonical adult inhibitory-control organization, this system is specifically compromised in children with ADHD. The striking spatial convergence between the core node recruited by TD children and the localized region of hypoactivation in the ADHD group identifies the preSMA as a critical site of neural dysfunction. Collectively, these results suggest that a reduced recruitment of a key cortical node in the inhibitory control network is associated with the disorder, offering a concrete neural target for developing novel intervention strategies.

## Data and code availability

The study-level data extracted for this meta-analysis (peak coordinates, sample sizes, tasks, and contrasts for each included experiment) and the analysis scripts will be made publicly available in an open repository upon publication of this article.

## Acknowledgements

This research was supported by National Institutes of Health MH124816 (W.C.), Stanford Psychiatry Innovator Grant (W.C.), and Stanford Maternal & Child Health Research Institute CE Grant (W.C.). The funders had no role in the study design, data analysis, the writing of manuscript, or the decision to submit it.

## Author contributions

Study design: W.C.; Data Extraction: E.K., S.P., H.H., W.C.; Data Analysis: E.K., S.P., H.H., Z.G., W.C.; Manuscript Drafting: E.K., W.C.; Manuscript Editing: E.K., S.P., H.H., Z.G., W.C.

