## Supplementary Material for "Functional deficits of the inhibitory control system in children with ADHD: A task-based fMRI meta-analysis"

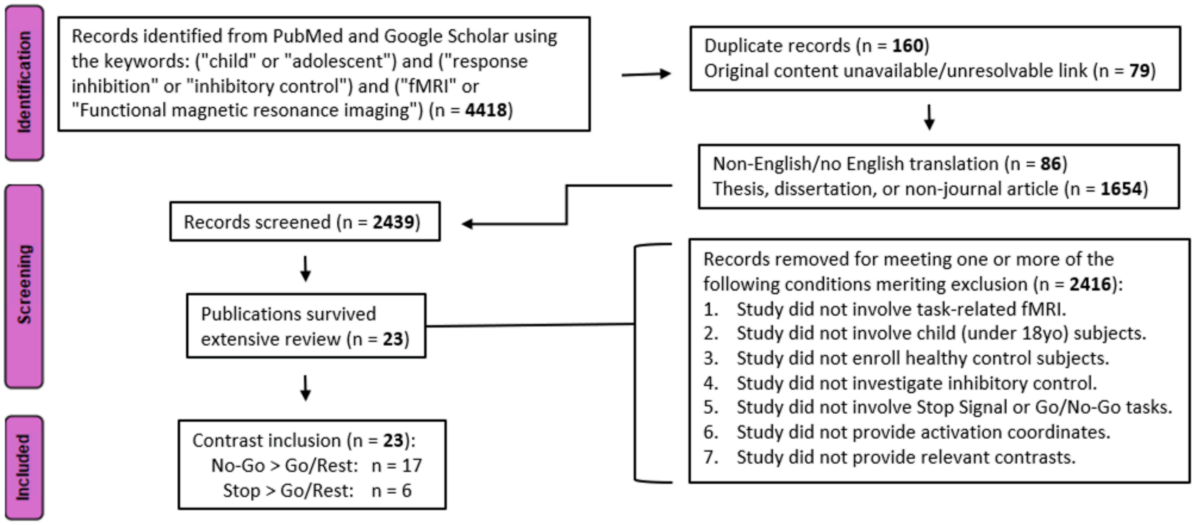


**Figure S1.** Flow chart of study identification and screening (TD children).

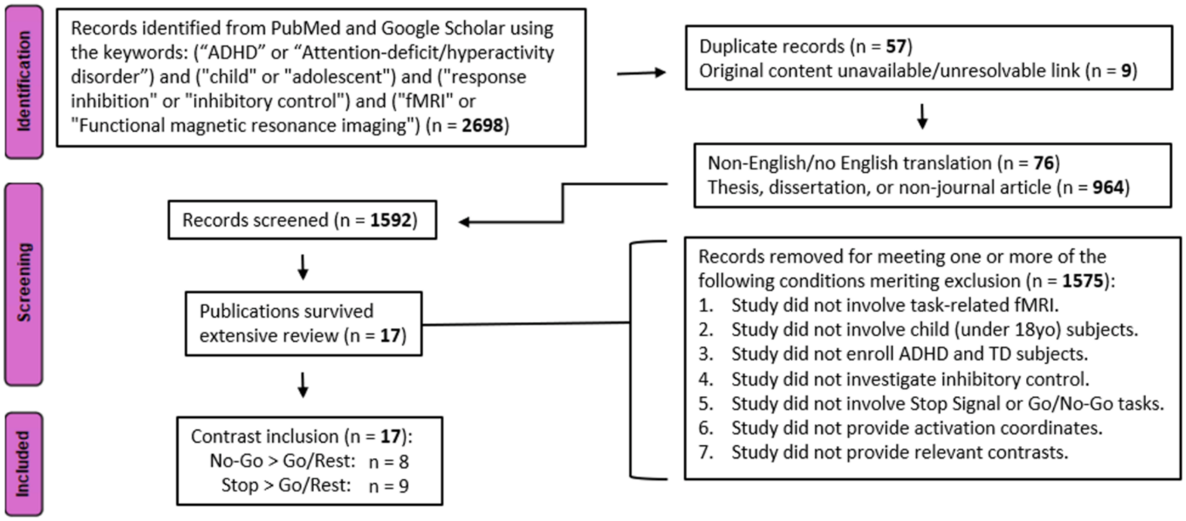


**Figure S2.** Flow chart of study identification and screening (TD children versus children with ADHD).

**Table S1.** List of fMRI studies of inhibitory control included in the meta-analysis of TD children.

| Year | First Author | Journal | Task | MRI Model | Contrast | Coordinate Space | Foci (n) | Mean Age | TD Sample (m/f) |
|---|---|---|---|---|---|---|---|---|---|
| 2002 | Bunge | *Neuron* | Modified GNG | 3T GE Signa | NoGo > Go | Talairach | 2 | 10 | 16 (9m/7f) |
| 2002 | Durston | *Developmental Science* | GNG | 1.5T GE Signa | NoGo > Go | Talairach | 3 | 8.7 | 10 (5m/5f) |
| 2003 | Booth | *NeuroImage* | Modified GNG | 1.5T GE (unspecified) | NoGo > Go | MNI | 16 | 10.9 | 12 (7m/5f) |
| 2003 | Durston | *Biological Psychiatry* | GNG | 1.5T GE Signa | NoGo > Go | Talairach | 7 | 8.68 | 14 (12m/2f) |
| 2004 | Schulz | *The American Journal of Psychiatry* | GNG | 1.5T GE Horizon | NoGo > Go | Talairach | 5 | 17.5 | 9 (9m/0f) |
| 2004 | Tamm | *Journal of the American Academy of Child and Adolescent Psychiatry* | Modified GNG | 1.5T GE Signa | NoGo > Go | Talairach | 3 | 15.58 | 12 (12m/0f) |
| 2005 | Booth | *The Journal of Child Psychology and Psychiatry* | Modified GNG | 1.5T GE (unspecified) | NoGo > Go | MNI | 16 | 10.9 | 12 (7m/5f) |
| 2006 | Durston | *Biological Psychiatry* | GNG | 1.5T Philips Gyroscan | NoGo > Go | Talairach | 9 | 15.27 | 11 (11m/0f) |
| 2006 | Pliszka | *The American Journal of Psychiatry* | SST | 2T GE Prestige | Stop > Rest | Talairach | 2 | 13.2 | 15 (9m/6f) |
| 2007 | Simmonds | *Neuropsychologia* | GNG | 1.5T Philips Powertrack | NoGo > Go | Talairach | 10 | 10.6 | 30 (13m/17f) |
| 2008 | Suskauer | *Journal of Cognitive Neuroscience* | GNG | 1.5T Philips Powertrack | NoGo > Rest | Talairach | 7 | 10.8 | 25 (15m/10f) |
| 2009 | Bennett | *Neurotoxicology and Teratology* | GNG | 1.5T GE (unspecified) | NoGo > Go | Talairach | 8 | 12 | 11 (5m/6f) |
| 2009 | Sheinkopf | *Developmental Neuroscience* | GNG | 1.5T Siemens (unspecified) | NoGo > Go | Talairach | 11 | 8.5 | 12 (7m/5f) |
| 2010 | Passarotti | *Psychiatry Research: Neuroimaging* | Modified SST | 3T GE Signa | Stop > Go | Talairach | 5 | 14.13 | 15 (7m/8f) |
| 2012 | Siniatchkin | *Brain Topography* | GNG | 3T Philips Achieva | NoGo > Go | MNI | 7 | 9.8 | 12 (9m/3f) |
| 2013 | Wang | *Clinical Neurophysiology* | GNG | 1.5T Siemens MAGNETOM Avanto | NoGo > Rest | Talairach | 1 | 10.23 | 31 (16m/15f) |
| 2014 | Bhaijiwala | *Brain and Behavior* | SST | 1.5T GE LX | Stop > Go | Talairach | 7 | 15.4 | 12 (unspec.) |
| 2014 | Cubillo | *Cerebral Cortex* | SST | 3T GE Signa | Stop > Go | Talairach | 9 | 13.75 | 29 (29m/0f) |
| 2014 | Heitzeg | *Drug and Alcohol Dependence* | GNG | 3T GE Signa | NoGo > Go | MNI | 6 | 10.9 | 19 (15m/4f) |
| 2015 | Janssen | *Psychiatry Research: Neuroimaging* | Modified SST | 1.5T Siemens Sonata | Stop > Stop-control | Talairach | 4 | 10.28 | 17 (13m/4f) |
| 2017 | van Hulst | *European Child & Adolescent Psychiatry* | Modified GNG | 3T Philips Achieva | NoGo > Go | MNI | 13 | 10.5 | 26 (26m/0f) |
| 2018 | Gooskens | *Developmental Cognitive Neuroscience* | SST | 3T Siemens Trio & Prisma/ 3T GE MR750/ 3T Philips Achieva | Stop > Rest | MNI | 11 | 10 | 53 (29m/24f) |
| 2021 | Cai | *Molecular Psychiatry* | GNG | 3T GE Signa | NoGo > Rest | MNI | 75 | 13.65 | 30 (22m/8f) |

**Table S2.** List of fMRI studies of inhibitory control included in the between-group meta-analysis (TD children vs. children with ADHD).

| Year | First author | Journal | Task | Diagnostic Method | ADHD subtype | MRI Model | Medication status | Contrasts | Coordinate Space | Foci (n) | Mean Age | Sample size (m/f) [TD/ADHD] |
|---|---|---|---|---|---|---|---|---|---|---|---|---|
| 1999 | Rubia | *American Journal of Psychiatry* | SST | Meeting DSM-IV Criteria for ADHD, Parental Account of Child Symptoms, Conners teacher rating scale | Hyperactive | 3T GE Signa | Unmedicated or 1-week washout prior to testing | TD > ADHD: Stop > Rest | Talairach | 4 | 14.71 | 16 (16m/0f) [9/7] |
| 2004 | Tamm | *Journal of the American Academy of Child and Adolescent Psychiatry* | Modified GNG | Diagnostic Interview for Children and Adolescents, Connors ADHD/DSM-IV Scale | Combined | 1.5T GE Signa | Unmedicated or 18-hour stimulant medication washout prior to testing | TD > ADHD: ADHD > TD: NoGo > Go | Talairach | 1 ; 1 | 15.81 | 22 (22m/0f) [12/10] |
| 2005 | Booth | *The Journal of Child Psychology and Psychiatry* | Modified GNG | Diagnosis by medical professional, Disruptive Behavior Rating Scale to confirm diagnosis | Combined/Unspecified | 1.5T GE Signa | 48-hour stimulant medication washout prior to behavioral testing or MRI scan | TD > ADHD: NoGo > Go | Talairach | 17 | 10.95 | 24 (15m/9f) [12/12] |
| 2006 | Durston | *Biological Psychiatry* | GNG | Diagnostic Interview Schedule for Children, DSM-IV criteria, Child Behavior Checklist | 10 Combined, 1 Hyperactive | 1.5T GE Horizon | Unmedicated or 24-hour stimulant medication washout prior to testing | TD > ADHD: NoGo > Go | Talairach | 5 | 14.62 | 22 (22m/0f) [11/11] |
| 2008 | Suskauer | *Journal of Cognitive Neuroscience* | GNG | Diagnostic Interview for Children and Adolescents - Fourth edition, Conners' Parent and Teacher Rating Scales - Revised | 17 Combined, 6 Inattentive, 2 Hyperactive/Impulsive | 1.5T GE Signa | Unmedicated or completed washout for stimulant medication day before and day of scan; No additional psychoactive medications allowed | TD > ADHD: ADHD > TD; NoGo > Rest | Talairach | 13 ; 1 | 10.8 | 50 (30m/20f) [25/25] |
| 2009 | Cerullo | *Early Intervention in Psychiatry* | SST | DSM-IV criteria | Unspecified | 1.5T GE (unspecified) | Unmedicated or withheld stimulant medication morning of scan | TD > ADHD; ADHD > TD; Stop > Rest | MNI | 1 ; 1 | 14.25 | 23 (14m/9f) [13/10] |
| 2010 | Passarotti | *Psychiatry Research: Neuroimaging* | Modified SST | Washington University Schedule for Affective Disorders and Schizophrenia, DSM-IV Criteria, Parent ADHD Rating Scale IV-Revised | Unspecified | 1.5T Philips Gyroscan | Unmedicated or gradual reduction of medication (mood stabilizers, second generation anti-psychotics, stimulants) over a 3-week period such that patients were drug-free for minimum of 7 days before testing | TD > ADHD: ADHD > TD; Stop > Go | Talairach | 7 ; 3 | 13.69 | 26 (13m/13f) [15/11] |
| 2011 | Braet | *Neuropsychologie* | Modified GNG | DSM-IV Criteria for ADHD, Child and Adolescent Psychiatric Assessment or Parental Account of Children's Symptoms | Unspecified | 2T GE Prestige | Unmedicated or 24-hour stimulant medication washout prior to testing | TD > ADHD: NoGo > Rest | Talairach | 2 | 13.55 | 58 (48m/10f) [38/20] |
| 2011 | Rubia | *Biological Psychiatry* | SST | Interviews with child psychiatrist using Maudsley Diagnostic Interview, DSM-IV Criteria, Strengths and Difficulties Questionnaire for Parents | Combined | 1.5T Philips Powertrack | Scanned twice 1 week apart, once after .3mg/kg of methylphenidate administration and once after placebo (vitamin C, 100 mg); No previous exposure to stimulant medication as recruitment criteria | TD > ADHD: Stop > Go | Talairach | 13 | 13 | 25 (25m/0f) [13/12] |
| 2012 | Siniatchkin | *Brain Topography* | GNG | Diagnosis by experienced neuropediatrician and child/adolescent psychiatrist, CBCL, German adaptive Diagnostic Checklist (DCL-HKS), DSM-IV Criteria | Combined, Hyperactive-Impulsive | 1.5T GE (unspecified) | Unmedicated or 24-hour stimulant medication washout prior to testing | TD > ADHD: NoGo > Go | Talairach | 3 | 9.65 | 24 (19m/5f) [12/12] |
| 2013 | Wang | *Clinical Neurophysiology* | GNG | Diagnosed based on DSM-IV criteria after clinical examination at outpatient Child Psychiatry Clinic, KSADS-PL, Conners' Parent Symptom Questionnaire | Combined | 1.5T Siemens (unspecified) | All participants stimulant medication-naive; 24-hour washout period for caffeine or other activity-level influencing foods or liquids | TD > ADHD; ADHD > TD; NoGo > Rest | Talairach | 6 ; 7 | 9.93 | 59 (41m/18f) [31/28] |
| 2014 | Bhaijiwala | *Brain and Behavior* | SST | PICS-IV, DSM-IV Criteria for ADHD based on parent, teacher and patient self-report, Global Assessment Scale | 9 Combined, 3 Inattentive | 3T GE Signa | Unmedicated or 24-hour stimulant medication washout prior to testing; Excluded if taking medication other than stimulants such as SSRI's and atomoxetine | TD > ADHD: ADHD > TD; Stop > Go | Talairach | 3 ; 2 | 14.6 | 24 (unspec.) [12/12] |
| 2014 | Cubillo | *Cerebral Cortex* | SST | Maudsley Diagnostic Interview, Strengths and Difficulties Questionnaire, Conners' Parent Rating Scale | Combined | 3T Philips Achieva | Randomized medication assignment for ADHD participants across three MRI Scans – Placebo (Vitamin C, 50 mg), MPX (Equasym, 0.3 mg/kg, range 5–20 mg), or ATX (Strattera, 1 mg/kg, range 16–66 mg). Participants remained medication free between scans, and medication was administered 1.5 hours before scanning. Control participants scanned once, unmedicated | TD > ADHD: ADHD > TD; Stop > Go | MNI | 4 ; 2 | 13.41 | 48 (48m/0f) [29/19] |
| 2014 | Hart | *Human Brain Mapping* | SST | Maudsley Diagnostic Interview, Strengths and Difficulties Questionnaire, Conners' Parent Rating Scale, Social Communication Questionnaire | Combined | 1.5T Siemens MAGNETOM Avanto | Unmedicated or 48-hour stimulant medication washout period prior to testing | TD > ADHD: ADHD > TD; Stop > Go | Talairach | 8 ; 1 | 14 | 60 (60m/0f) [30/30] |
| 2015 | Janssen | *Psychiatry Research: Neuroimaging* | Modified SST | Diagnosis from child psychiatrist at outpatient mental health facility based on DSM-IV criteria, Diagnostic Interview Schedule for Children, Disruptive Behavior Disorders Rating Scale | 19 Combined, 2 Inattentive | 1.5T GE LX | Unmedicated or 48-hour stimulant medication washout period prior to testing | TD > ADHD: ADHD > TD; Stop > Stop-control | Talairach | 3 ; 2 | 10.45 | 38 (32m/6f) [17/21] |
| 2015 | van Rooij | *American Journal of Psychiatry* | SST | Schedule for Affective Disorders and Schizophrenia for School-Age Children, Conners' Rating Scales | Unspecified | 3T GE Signa | Unmedicated or 48-hour psychoactive medication washout period prior to testing | TD > ADHD: Stop > Go | Talairach | 5 | 16.9 | 309 (184m/125f) [124/185] |
| 2018 | Stevens | *Biological Psychiatry: Cognitive Neuroscience and Neuroimaging* | GNG | Schedule for Affective Disorders and Schizophrenia for School-Age Children–Present and Lifetime version | Combined | 1.5T Siemens Sonata | Unmedicated or 24-hour stimulant medication washout prior to cognitive and MRI assessments | TD > ADHD: ADHD > TD; NoGo > Rest | Talairach | 19 ; 1 | 15 | 85 (unspec.) [63/22] |
